# The rate of ecosystem acclimation is the dominant uncertainty in long-term projections of an ecosystem service

**DOI:** 10.1101/2021.08.11.455579

**Authors:** Andrew J. Felton, Robert K. Shriver, Michael Stemkovski, John B. Bradford, Katharine N. Suding, Peter B. Adler

## Abstract

Rapid climate change may exceed ecosystems’ capacity to respond through processes including phenotypic plasticity, compositional turnover and evolutionary adaption. However, research predicting impacts of climate change on ecosystem services rarely consider this rate of “ecosystem acclimation.” Combining statistical models fit to historical climate data and remotely-sensed estimates of herbaceous productivity with an ensemble of climate models, we demonstrate that assumptions concerning acclimation rates are a dominant source of uncertainty: models assuming minimal acclimation project widespread decreases in forage production in the western US by 2100, while models assuming that acclimation keeps pace with climate change project widespread forage increases. Uncertainty related to ecosystem acclimation is larger than uncertainties from variation among climate models or emissions pathways. A better understanding of ecosystem acclimation is essential to improve long-term forecasts of ecosystem services, and shows that management to facilitate ecosystem acclimation may be necessary to maintain ecosystem services at historical baselines.

## Introduction

Anthropogenic climate change is disrupting relationships between climate and ecosystem structure and function that have developed over millennia (Whittaker 1975; Svenning & Sandel 2013). Ecosystems are responding to this disruption through mechanisms spanning a wide range of timescales (Smith *et al*. 2009). Individual physiological rates can change on scales of minutes, population sizes typically change over years to decades, and species turnover, evolutionary adaptation, and alteration of disturbance regimes and biogeochemical cycles may play out over decades or centuries (Fig. 1). While the term “acclimation” has traditionally referred to phenotypic plasticity, here we consider “ecosystem acclimation” as the net effect of all the processes by which ecosystems respond to climate change. Given the long timescales of many of these processes, ecosystem acclimation will likely lag behind the pace of climate change, limiting the rate of ecosystem acclimation and creating disequilibria between ecosystem structure and climate (Webb 1986; Svenning & Sandel 2013; Blonder *et al*. 2015; Williams *et al*. 2021). The consequences of such disequilibria for ecosystem functioning remain unknown.

**Figure 1.**
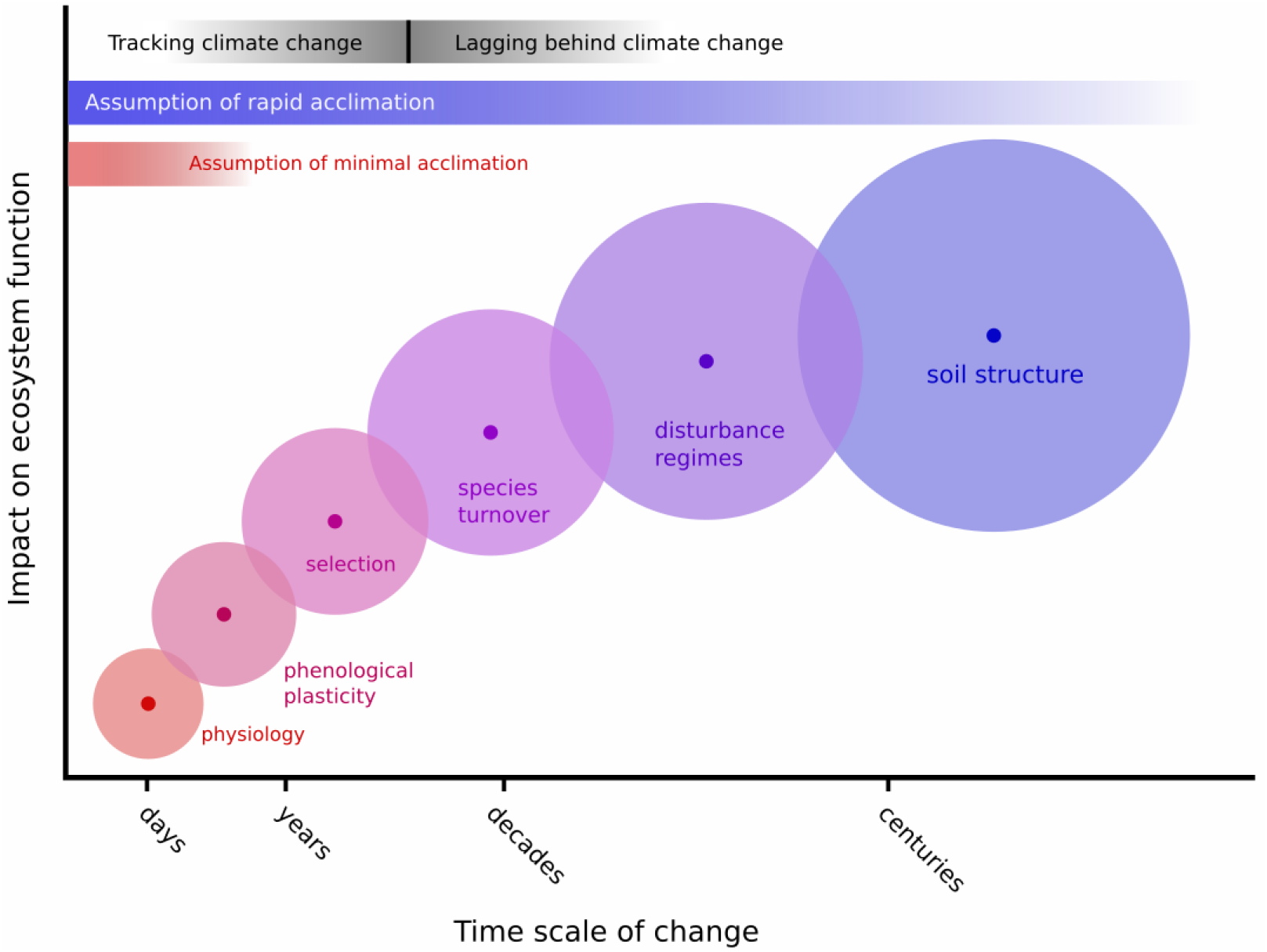
“Ecosystem acclimation” refers to the many processes by which ecosystems respond to climate change. The fastest processes, such as physiological adjustment or phenotypic plasticity, may keep pace with the rapidly changing climate. Slower processes, such as selection for drought-tolerant genotypes or species, species colonization and extinction, or changes in disturbances regimes and soil structure in terrestrial systems, may lag behind the changing climate, leading to loss of ecosystem services. We assume that the slowest processes will have the greatest impacts on ecosystem functioning in the long-term (a grassland cannot turn into a desert without species turnover and changes in disturbance regimes and soils), but that the uncertainty surrounding these impacts (size of circles) is also greater because dynamics of slow processes are harder to study. Models to project ecological impacts of climate change often make extreme, but implicit, assumptions about which processes will influence the focal ecological response (red and blue horizontal bars).

Our lack of understanding about the timescales and potential impacts of acclimation lags (Fig. 1) creates great uncertainty in long-term projections of climate change impacts on ecosystem services (Carpenter 2002; Luo *et al*. 2011; Charney *et al*. 2016; Runting *et al*. 2017). While the fast components of ecosystem acclimation, such as physiology, can be studied using experiments and observations, we know much less about the slow components. Experiments typically lack the spatial or temporal extent to observe species turnover or evolutionary adaptation, and may only capture transient behavior (Collins *et al*. 2012; Reich *et al*. 2018). Paleoecological observations provide much of what we know about long-term ecological responses to environmental change (Jackson & Overpeck 2000), but the data have critical gaps and current rates and magnitudes of warming exceed those observed in the past (Kaufman *et al*. 2020). Without relevant empirical data about the slow components of acclimation, most ecological models assume either minimal ecosystem acclimation or rates of acclimation that perfectly track climate change (Blonder *et al*. 2017). Either way, the sensitivity of results to these assumptions is rarely tested. Although uncertainty about the rate of ecosystem acclimation has been recognized (Adler *et al*. 2020; Luo & Schuur 2020; Rollinson *et al*. 2021), to our knowledge previous studies have not estimated its magnitude or consequences for ecosystem services.

Here, we quantify how varying assumptions about the rate of ecosystem acclimation translates into uncertainty in long-term projections of an economically important ecosystem service, forage production. We compared this uncertainty to those from more commonly studied sources, such as variation in greenhouse gas emission scenarios (referred to as relative concentration pathways or RCPs), general circulation models (GCMs), spatial variation, and model parameter error. Our approach exploits differences in how forage production responds to spatial and temporal variation in climate (Lauenroth & Sala 1992). Spatial relationships between forage production and climate capture an advanced stage of ecosystem acclimation: species composition, disturbance regimes, and resource pools have had centuries to equilibrate to a relatively stationary climate. When we project future production based on spatial gradients, we assume that even the slow components of ecosystem acclimation instantaneously track the rate of climate change, representing a scenario of rapid ecosystem acclimation (Fig. 1). In contrast, when we study the relationship between historical, interannual variation in weather and interannual variation in production at one location, we hold slow processes such as changes in species composition and resource supply relatively constant (Briggs & Knapp 1995). Projections of future forage production based on this time-series approach represent a scenario of minimal ecosystem acclimation driven only by mechanisms operating on fine timescales (Fig. 1). Together, the spatial gradient, “rapid acclimation” scenario and the time-series, “minimal acclimation” scenario provide objective bounds on the range of uncertainty in future forage production resulting from different assumptions about the rate of ecosystem acclimation.

We implemented this approach by fitting statistical models to 30 years of remotely-sensed estimates of forage production (Robinson *et al*. 2019) for six rangeland ecoregions covering the western U.S. (Fig. S1). These data allowed us to simultaneously describe how productivity responds to spatial gradients in climate and temporal variation in weather by decomposing precipitation and temperature observations into spatial and temporal components. The spatial component is represented by variation in mean precipitation and temperature across sites (pixels), while the temporal component is represented by annual anomalies from the mean precipitation and temperature at each site. Because our statistical model contained both spatial and temporal relationships between climate and productivity, we were able to use either the time-series (minimal acclimation) or spatial gradient (rapid acclimation) approach to project effects of climate change on forage production (Fig. S2).

## Material and Methods

### Rangeland ecoregions

We studied five expansive rangeland ecoregions of the western United States: the Northern Mixed Prairies (NM), the Shortgrass Steppe (SGS), the Cold Deserts (CD), the California Annual grasslands (CA), and the Hot Deserts (a combination of the Mojave, Sonoran, and Chihuahuan Deserts). These ecoregions were defined by potential natural vegetation (Kuchler 1964). A previous assessment of functional type cover in each of these ecoregions was consistent with expectations based on potential natural vegetation (Felton *et al*. 2021). We split the Hot Deserts ecoregion into two to account for differences in precipitation seasonality. We assigned sites receiving > 30% of total annual precipitation during the summer months (July-September) to a Hot Desert summer precipitation (HDS) ecoregion, and assigned sites receiving less than 30% of annual precipitation during summer to a Hot Desert winter precipitation ecoregion (HDW) (Fig. S1).

### Forage production data

We leveraged a remotely-sensed dataset of annual net primary productivity data developed specifically for rangeland ecosystems of the western US, spanning the years 1986-2015 (Robinson *et al*. 2019). This data product partitions annual primary production (defined as net carbon inputs in gm^-2^) into functional groups of shrubs, trees, annual forbs/grasses, and perennial forbs and grasses. We defined annual forage production, our response variable, as the sum of productivity from the herbaceous functional groups: annual and perennial grasses and forbs. We aggregated these data from their native spatial resolution (30 m) to 1/16^th^ degree (approximately 6.9 km) to match the spatial resolution of our climate data.

We applied a multi-step filtering process to remove non-rangeland pixels, such as irrigated agriculture (Felton *et al*. 2021). We removed 6.9 km pixels in which less than 50% of the 30 m LANDSAT subpixels were rangeland. We also removed pixels where the average net primary productivity was below 10 gm^-2^. We then sequentially removed outlier pixels for each ecoregion based on distributions of multiple metrics of plant-water relationships across each ecoregion. These metrics, presented in order, were 1) mean precipitation use efficiency (mean net primary productivity/mean annual precipitation), the slope of annual productivity regressed on annual precipitation, and 3) annual precipitation use efficiency. Outliers were defined as beyond three standard deviations from the mean. After filtering, we had 774 pixels in California Annual grasslands, 13,869 in Cold Desert, 6,446 in Hot Deserts summer precipitation, 1,573 in Hot Deserts winter precipitation, 13,070 in the Northern Mixed Prairie, and 4,925 in the Shortgrass Steppe.

### Historical climate data and model selection

For historical climate data, we used a gridded hydrometeorological data product with daily time steps of temperature and precipitation (Livneh *et al*. 2013, 2015). The native resolution of these data are ∼6 km, which we aggregated to 6.9 km to match the resolution of soil moisture data used in our pre-model fitting. These data encompass years 1915-2015, but we focused on years 1986-2015 to correspond with the temporal coverage of the forage production data.

Explaining an annual ecological response, like production, using daily weather data poses a difficult model selection problem because of the virtually limitless choices about how to aggregate the weather data (Tredennick *et al*. 2021). Therefore, we constrained our covariate selection approach in two ways: First, in each candidate model we considered only one variable representing moisture stress and one variable representing heat stress, along with their interactions as explained below. This constraint guarded against collinearity and overfitting. In addition to precipitation (moisture stress) and temperature (heat stress), we considered additional covariates simulated by the SOILWAT ecohydrological model (Bradford *et al*. 2014; Schlaepfer & Andrews, C. A. 2018), using the daily weather data (Livneh *et al*. 2013) as input for each location (pixel). These included moisture-related variables such as soil water availability, volumetric water content, and transpiration, and variables related to heat stress such as potential evapotranspiration (PET) and the ratio of soil water availability to PET. Preliminary analyses using linear mixed effects models (not shown) indicated that precipitation and temperature explained more variation in observed productivity than pairs of variables derived from SOILWAT. For all subsequent models reported here, the raw covariates were precipitation and temperature.

Second, we constrained variable selection using our *a priori* knowledge of each ecoregion to define the most relevant climate window, rather than exhaustively searching for the optimal temporal window over which to aggregate the moisture stress and heat stress variables. For the California annual grasslands, the cold deserts, and the hot deserts with winter precipitation, we summed precipitation and averaged temperature over fall, winter, and spring (October – June). For the northern mixed prairies, shortgrass steppe, and hot deserts receiving summer precipitation, we aggregated over spring and summer (April – September).

While the raw covariates for each ecoregion were total precipitation and mean temperature aggregated over the relevant seasons, we decomposed each of these raw covariates into spatial means and temporal (annual) anomalies and included interactions among them (Kleinhesselink & Adler 2018; Felton *et al*. 2021). Specifically, we decomposed *P*_*i,t*_, the precipitation received in pixel *i* and year *t*, into the mean precipitation in pixel 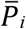, and the annual precipitation anomaly in pixel *i* and year *t, δP*_*i,t*_. Note that 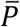 varies only across space while δP varies only in time. Similarly, we define 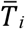 as the mean temperature in pixel *i* and *δT*_*i,t*_ as the annual temperature anomaly. We then designed interactions among these derived covariates to allow the effects of annual anomalies to vary in space, as shown in Table S1. (Note that in our computer code, we refer to the precipitation and temperature anomalies as deviations.)

### Model fitting

We developed spatiotemporal hierarchical models for each ecoregion to infer the response of forage production (herbaceous NPP) to past climate conditions, and then used this model to predict changes in forage in response to future climate change.

Observed forage production, *y*, in each pixel, *i*, and year, *t*, was modeled as normally distributed with mean (*µ*_*i,t*_) and process variance 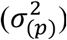:

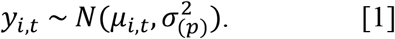

The process model for the mean is a linear model of covariates **X**_*i,t*_ and ecoregion-specific parameters **b**, along with additive, pixel-specific spatial (*η*_*i*_) and year-specific temporal (*τ*_*t*_) random effects:

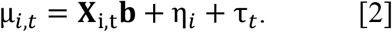

Year random effects were fit with a non-centered parameterization (more computationally efficient in Stan) using a standard normal distribution, rescaled with standard deviation *σ*_(*t*)_:

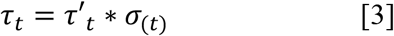

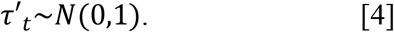

Spatial random effects were fit using a spatial dimension reduction approach following the general methods of (Tredennick *et al*. 2016). Dimension reduction approaches help overcome the computational burden of modeling residual autocorrelation structures by reducing the number of pixels to a constituent number of knots that summarize the landscape of spatially autocorrelated processes not accounted for by covariates. Spatial random effects were modeled as

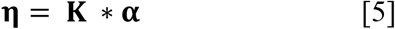

where **η** is a vector of spatial random effects specific to each pixel, *i* (i = 1…*I*), **α** is a vector of spatial random effects specific to each knot, *s* (*s* = 1…*S*), and **K** is an *I* by *S* matrix that describes how the knot based random effects are translated into random effects for each pixel.

Each element in **K**, *K*_*i,s*_, is a normalized, distance weighted function from each knot to pixel

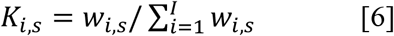

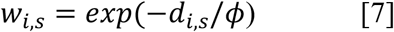

where *d*_*i,s*_ is the distance from pixel *i* to knot *k*, and *ϕ* is a distance decay parameter.

Knot-based random effects were fit with a non-centered parameterization using a standard normal distribution, rescaled with standard deviation *σ*_(*s*)_ as:

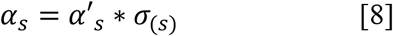

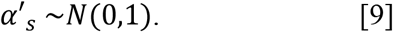

Following the approach of (Tredennick *et al*. 2016), *ϕ* and knot spacing was determined empirically prior to model fitting to improve computation based on the observed scale of spatial auto-correlation. To do this, we fit a version of our model including only fixed effects, using the maximum likelihood solution for **b**. We then calculated model residuals between our fixed-effects-only model and observed NPP values, and visualized the scale of autocorrelation using empirical semi-variograms (Fig. S3). The spacing of knots is determined visually by the approximate sill distance of each ecoregion, and value of *ϕ* is one-third the sill distance. Knots were placed in a regular grid across each ecoregion with spacing based on sill distance (Fig. S4).

To limit overfitting, we regularized covariates with Bayesian ridge regression. This was done by fitting seven different models in each ecoregion to a subset of the data (randomly withholding 5 years, ∼17% of the data) with increasingly wider zero centered priors for covariate parameters (**b∼*N*(**0, **I***υ*^2^)), and then testing the predictive ability of each prior specification with the withheld data (Hooten & Hefley 2019). The priors for **b** that minimized the average mean squared predictive error (MSPE) in each ecoregion were then used to fit the final model to the full dataset (Fig. S5). For each ecoregion, this final model was used to project future forage production. Parameter estimates are shown in Table S2, and goodness-of-fit measures are shown in Table S3.

### Future climate data

We focused our projections on the late-century time period, defined as 2061-2100. To represent the range of uncertainty in future climate data, daily climate data from eleven different general circulation models (GCMs) under two different representative concentration pathways (RCPs) families; scenario 4.5, which represents a relatively low emissions scenario, and scenario 8.5, which represents the highest emissions scenarios (Moss *et al*. 2010; Taylor *et al*. 2012), were used to drive the model. The GCMs were selected both to represent the diversity of model projections among GCMs included in CMIP5 (Knutti *et al*. 2013), and to include models that performed well in hindcast comparisons within the western U.S. (Rupp *et al*. 2013) (Rupp unpublished data). The 11 GCMs we used were: CESM1-CAM5, CSIRO-Mk3-6-0, CNRM-CM5, FGOALS-g2, FGOALS-s2, GISS-E2-R, HadGEM2-ES, inmcm4, IPSL-CM5A-MR, and MIROC-ESM. Future climate data was extracted as monthly time-series from 1/8° downscaled and bias-corrected products of the fifth phase of the Climate Model Intercomparison Project (CMIP5) (Maurer *et al*. 2007).

### Forage projections and uncertainty and sensitivity analyses

We combined the parameters estimates of the statistical model with GCM simulations of late century precipitation and temperature to project future forage production. The first step was decomposing future precipitation and temperature into means and annual anomalies for each location (pixel) (Fig. S2). One option is to calculate future annual anomalies in precipitation and temperature based on the *historical* means at each location. This implements a time-series approach: we assume that climate means will not change, but there will be many hotter than normal years in the future, and the effects of these anomalous years are captured by the model’s temporal anomaly terms. The second option is to calculate future annual anomalies based on the *future* means at each location. This implements the spatial gradient approach: we assume relatively small changes in the annual anomalies but potentially large changes in mean precipitation and temperature, and the effects of these shifts in the means are captured in the model’s spatial terms. Although most of our analyses focus on the time-series (historical means as the reference) and spatial gradient (future means as the reference) approaches, we also explored a third option in which we calculate annual anomalies based on GCM projections for the mid-century period (2021-2060). This approach represents an intermediate rate of ecosystem acclimation.

After calculating the climate covariates, we projected future productivity. For each location and each year in the late century period, we predicted productivity for 22 climate scenarios (11 GCMs × 2 RCPs), 500 sets of model parameters drawn from the MCMC chains, and our two methods (time-series and spatial gradient). We then averaged predicted productivity across years within each pixel, preserving variation among all combinations of locations, parameter draws, GCMs, RCPs, and projection method. Because we extrapolated linear relationships, for some locations our models project negative values of forage production, resulting in declines greater than 100%. We chose to report these biologically impossible values, rather than truncating them at zero, to avoid confusion about our approach.

The uncertainty analysis featured in the main text methods focuses on the future change in mean forage production relative to the historical period. We calculated change in a given pixel as the difference between the mean annual production in the future period and mean annual production from the historical period, divided by the historical mean. Note that the historical mean is not based on observed data, but rather on predictions from the statistical model, ensuring that biases in the model do not influence the estimated change in productivity. Dividing by the historical mean led to some very large values in pixels where the historical mean was close to zero. Our uncertainty analysis was sensitive to these outliers, so we removed sites with predicted historical mean production less than 10 g m^-2^ (26 out of 13,839 pixels in the Cold Deserts, 18 of 6,446 pixels in the Hot Deserts with summer precipitation, and 14 of 1,573 pixels in the Hot Deserts with winter precipitation). Because our predictions for the two periods assume the same model process error, process error cannot contribute to change in productivity and we can ignore it. Similarly, we can ignore random year effects, which we assumed have the same variance in the future as in the historical period.

To perform the calculations, we loaded all the projected changes in forage production into an array with the following dimensions: 1) sites, 2) parameters draws from the posterior distribution (500), 3) GCMs (11), 4) RCPs (2), 5) rate of acclimation (2). We then analyzed variation in this array using two approaches. First, we took a sums-of-squares approach, quantifying the proportion of the total sum-of-squares contributed by each of the five main effects (the dimensions of the array), as well as the four two-way interactions involving rate of acclimation. Second, we quantified the standard deviation of the marginal means of each main effect. For example, we calculated the mean change in productivity at each site, averaging over parameters, GCMs, RCPs, and rate of acclimation, then computed the standard deviation among site means as a measure of the variation among sites in projected change in forage production.

We repeated this analysis of standard deviation for variation in absolute forage production, rather than variation in the change in forage production. For this case, we also included variation due to process error and random year effects, as estimated by the statistical models.

Note that results of both the sum-of-squares partitioning and the analysis of standard deviation are affected by the number of levels of each main factor (e.g., sites, parameters, GCMs, RCPs, projection methods). Because we consider only the two extremes for the rate of ecosystem acclimation, our estimate of the relative contribution from this source represents an upper bound. New information on rates of acclimation could reduce this uncertainty either by moderating the extremes, or by working with objective distributions of rates of acclimation (as we currently do with parameter error).

We also conducted a sensitivity analysis to help us understand the projections. For each ecoregion, we used the point estimates of model parameters to quantify the expected change in production at each pixel caused by 1) adding 1 cm to all mean precipitation values, 2) adding 1 cm to all annual precipitation anomalies, 3) adding 1°C to all mean temperature values, and 4) adding 1°C to all temperature anomalies. These perturbations fall well within the range of expected future changes in precipitation and temperature (Fig. S6). We recalculated all interaction terms after each perturbation. We report the mean change across locations for each of these four perturbations.

We performed a final set of projections to represent an intermediate rate of ecosystem acclimation, using GCM output for mid-century as the reference for calculating each pixel’s climate means and annual anomalies for the late-century period. We did not repeat the full uncertainty partitioning, but simply compared the averaged projected change in forage production for each pixel to the average change projected by the time-series (historical climate reference period) and spatial gradient (late-century climate reference period) approaches.

## Results

The minimal acclimation and rapid acclimation scenarios led to dramatically different projections of forage production by late century (2061-2100) (Fig. 2). Models assuming minimal acclimation projected widespread declines in forage production across western US rangelands. The projected declines were greatest in the shortgrass steppe and the hot desert ecoregions. In contrast, models assuming rapid acclimation projected widespread increases in forage production or, at worst, modest decreases. Across 84% of the Cold Deserts and 89% of the Northern Mixed Prairies, projected changes switched from negative under the minimal acclimation assumption to positive under the rapid acclimation scenario. For the other ecoregions, the two approaches projected a consistent direction, if not magnitude, of change in forage production, with increases in California and decreases in the Hot Deserts and Shortgrass Steppe.

**Figure 2.**
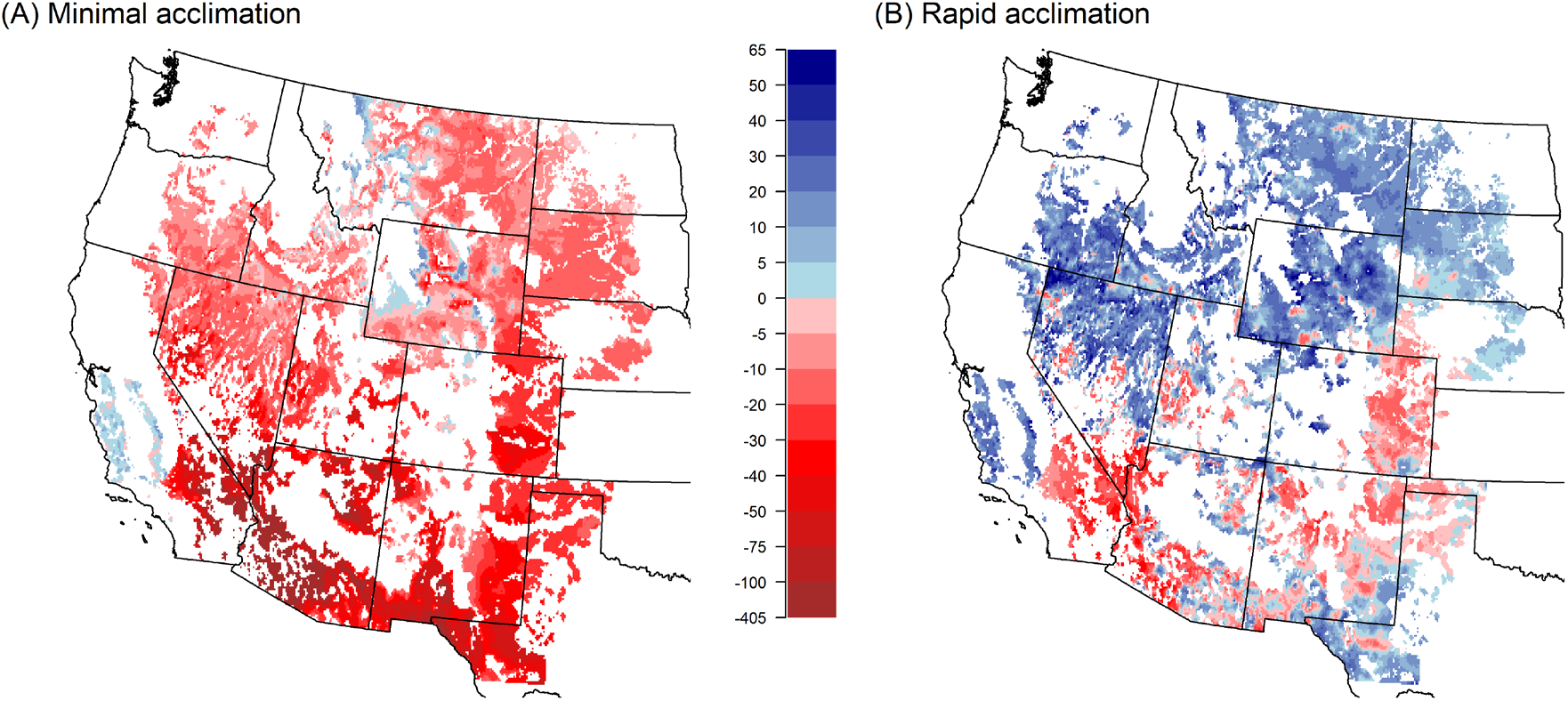
Projected changes in forage production during the late-century period differ between the (A) minimal and (B) rapid acclimation scenarios. Change is expressed as percentage of the historical baseline at each location. Values are means of projections based on 11 GCM ensemble members for the RCP 8.5 emissions pathway. Areas that are not natural rangeland vegetation were excluded from the analysis, and are shown in white.

A sensitivity analysis clarified why differences emerged between the minimal and rapid acclimation scenarios: positive annual temperature anomalies decreased forage production, but spatial variation in mean annual temperature had weak effects (Fig. 3). Essentially, production decreases during hot years but not necessarily in hot locations in all regions except California, which has a cool, winter-spring growing season. Because the minimal acclimation approach represents future temperature increases as temporal anomalies, it leads to strong negative effects of increasing temperatures, while the rapid acclimation approach represents future temperature increase as shifts in mean temperature, which have little impact on production. Temperature sensitivities estimated from field-based estimates of production at five sites indicate that these results are unlikely to be an artifact of the algorithm used to generate the herbaceous production product we used (SI Supporting Methods and Table S4).

**Figure 3.**
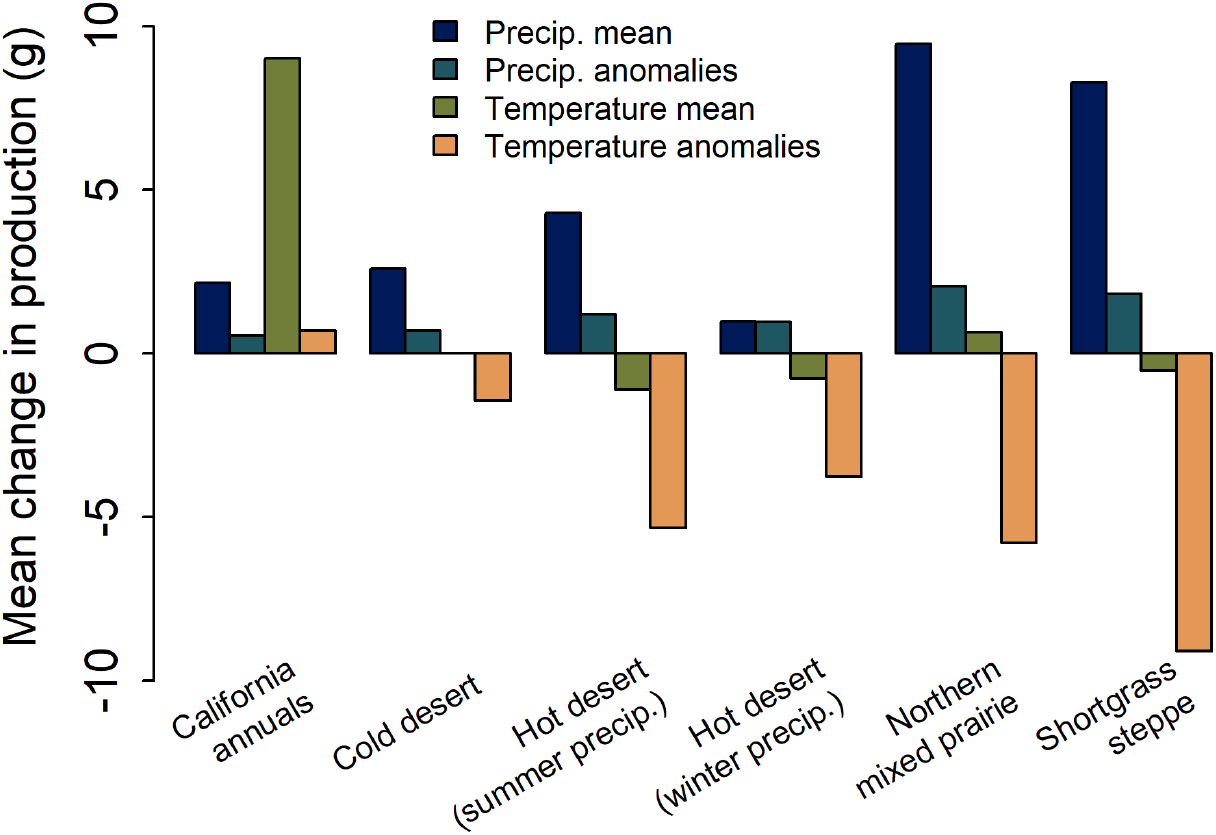
Forage production is most sensitive to temporal anomalies in temperature and spatial variation in mean precipitation. Bars show the change in production in late century projections, averaged across each ecoregion, in response to: 1) a 1 cm increase in mean annual precipitation, 2) a 1 cm increase in annual precipitation anomalies, 3) a 1°C increase in mean annual temperature, and 4) a 1°C increase in annual temperature anomalies. The temporal anomalies drive changes projected by the minimal acclimation scenario, while spatial variation drives changes projected by the rapid acclimation scenario.

Variation in mean annual precipitation had strong positive effects on production in all ecoregions, while precipitation anomalies had much weaker effects (Fig. 3). For example, predicted increases in precipitation (Fig. S6) in the Cold Deserts and Northern Mixed Prairies had strong positive effects on the rapid acclimation projections, whereas increasingly positive precipitation anomalies had little effect on projections assuming minimal acclimation.

The rate of ecosystem acclimation was the dominant source of uncertainty in projections for five of six ecoregions, contributing more than any other single source of uncertainty we considered (Fig. 4). The exception was the Hot Desert summer precipitation ecoregion, where spatial variation contributed most to uncertainty. The main effect of acclimation rate plus its two-way interactions with the other sources of uncertainty contributed from 44% (Northern Mixed Prairies) to 54% of the total variation (Hot Deserts with summer precipitation) (Table S5). Figs. S7 and S8 further emphasize that variation in projected change among GCMs and RCPs is small relative to the difference between the minimal and rapid acclimation scenarios.

**Figure 4.**
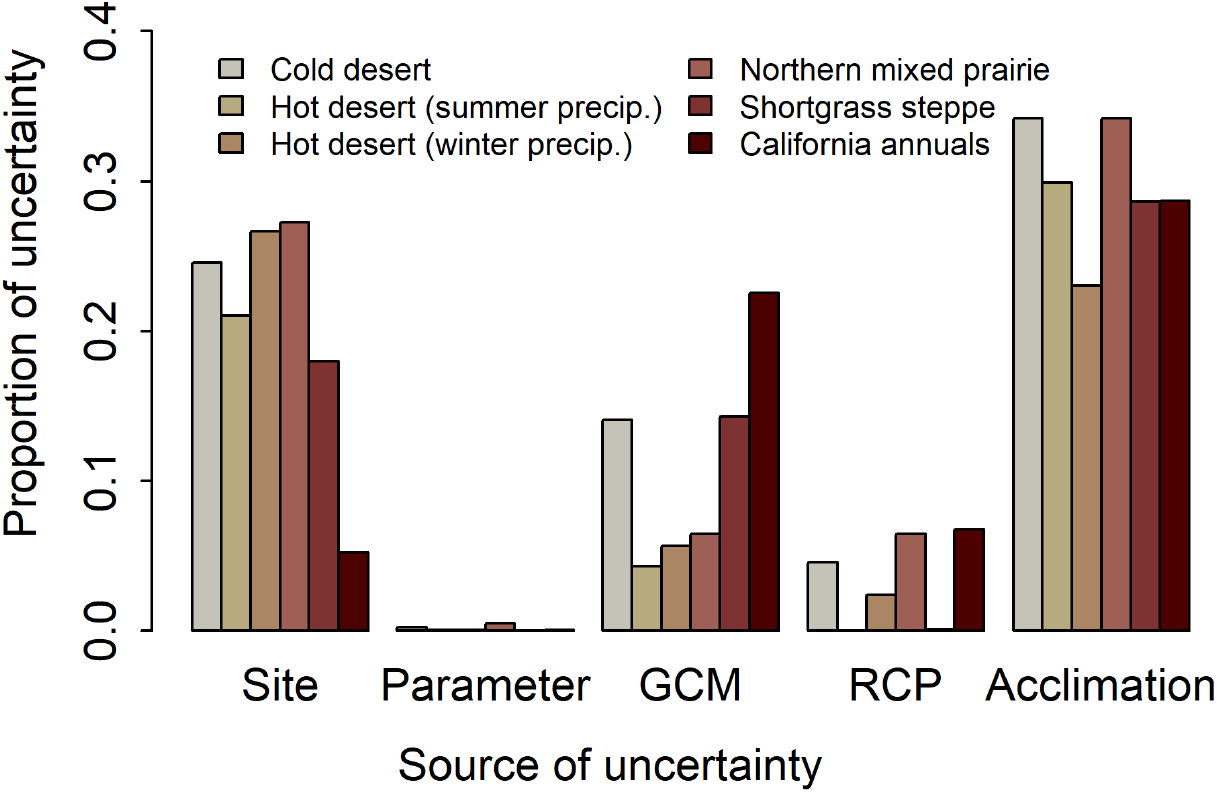
In five of six ecoregions, the rate of ecosystem acclimation is the largest source of uncertainty in projections of future changes in forage production. Uncertainty was partitioned as the proportion of the total sum-of-squares of projected changes in production by late-century. Site: spatial variation among pixels with each ecoregion. Parameter: variation among model parameter estimates in the posterior distribution. GCM: variation among global circulation models. RCP: variation among greenhouse gas emission scenarios. Acclimation: variation between the minimal and rapid ecosystem acclimation scenarios.

Could ecosystem acclimation help maintain historical levels of forage production? In the Hot Deserts and Shortgrass Steppe, where GCMs predict decreasing precipitation, our models project production declines even under rapid acclimation. In California, they predict production increases even under minimal ecosystem acclimation. For many locations in the Cold Deserts and Northern Mixed Prairies, however, the models predict a switch from decreased production under minimal acclimation to increased production under rapid acclimation. We also considered an intermediate rate of acclimation by using mid-century GCM projections (2021-2060) as the reference for calculating late-century annual anomalies. Median changes in production under this intermediate acclimation scenario were close to zero in the Cold Deserts but were still negative in the Northern Mixed Prairie (Fig. S9).

## Discussion

Our most important result is that much of the uncertainty in future forage production comes from the unknown rate of ecosystem acclimation. In fact, uncertainty about acclimation was much larger than uncertainties about future climate conditions. While studies of ecological impacts of climate change often include projections from an ensemble of climate models, or consider multiple emission scenarios, they rarely consider ecosystem acclimation as a potential source of uncertainty because assumptions about rates of acclimation are typically implicit in ecological models (Blonder *et al*. 2017). As a result, this source of uncertainty is often overlooked entirely. A better understanding of ecosystem acclimation could greatly reduce uncertainty in long-term projections of ecosystem services, independent of progress in climate science. We do not need precision; even estimates of acclimation rates and their ecosystem impacts (Fig, 1) to the nearest order of magnitude could greatly reduce the uncertainty of projections such as ours by identifying the appropriate climate reference period, or by informing a model weighting approach (Adler *et al*. 2020). Comparing simple models fit to time-series and spatial gradient data (Wilcox *et al*. 2016; Amburgey *et al*. 2018; Kleinhesselink & Adler 2018; Klesse *et al*. 2020) is a valuable first step in quantifying uncertainty related to ecosystem acclimation.

Our sensitivity analysis (Fig. 3) suggests fundamentally different patterns of acclimation to future changes in temperature and precipitation. In most ecoregions, production was far more sensitive to annual temperature anomalies than to spatial variation in mean temperatures. Because the minimal acclimation scenario estimates temperature effects based on annual anomalies, it projected much stronger negative impacts of future temperature increases compared to the rapid acclimation scenario, in which temperature effects reflect spatial gradients in climate and production. The differing effects of temperature means and anomalies are consistent with a recent study of tree growth (Klesse *et al*. 2020) and suggest that at any given location an unusually hot year can limit production either by creating heat stress (Breshears *et al*. 2021) or exacerbating water stress (Hoover *et al*. 2014; Williams *et al*. 2020). In contrast, ecosystems have adapted to spatial variation in mean temperatures through a variety of mechanisms, including shifts in phenology (Rice *et al*. 1992) or species composition (Edwards & Still 2008). We found the opposite pattern for precipitation: production was much more sensitive to spatial variation in mean precipitation than to temporal variation in annual precipitation anomalies, consistent with previous work (Huxman *et al*. 2004; Felton *et al*. 2021). Thus, the positive effects of future increases in precipitation, predicted for the Northern Mixed Prairies, have a much stronger influence under the rapid acclimation approach than for the minimal acclimation scenario. Overall, these results suggest that acclimation processes may help dryland ecosystems maintain production in the face of long-term temperature increases (e.g., Liu *et al*. 2018), but have less potential to limit impacts of changes in mean precipitation.

Could process-based models, an alternative to the phenomenological statistical models we built, accurately capture the mechanisms driving ecosystem acclimation? While possible in principle, we are skeptical that current process-based approaches could solve this problem. Process-based ecosystem models are often forced to make the same choice between time-series and spatial gradient approaches: studies focused on reproducing ecosystem dynamics typically tune their parameters using longitudinal data (e.g., Rollinson *et al*. 2017) while studies seeking to reproduce the broad-scale distribution of biomes tune their parameters with spatial data (e.g., Fisher *et al*. 2015). This choice represents an unrecognized assumption about the rate of acclimation. Furthermore, it may be computationally infeasible to simulate these models over the long temporal and broad spatial scales of our analysis, especially with the taxonomic or functional resolution necessary to capture the compositional changes central to ecosystem acclimation.

The second important result from our analysis is that rapid acclimation may be required to maintain ecosystem services at historical levels in most western US rangelands. Unfortunately, such rapid acclimation may be unlikely in dryland ecosystems. A decade or more of experimental precipitation reduction or addition appears necessary to significantly alter abundances of dominant plant species (Evans *et al*. 2011; Collins *et al*. 2012; Estiarte *et al*. 2016). Paleoecological studies document dramatic compositional changes in the recent past, but cannot provide sub-century resolution (Mottl *et al*. 2021). Changes in soil structure and soil resources will also be important, and likely even slower to change (Burke *et al*. 1995). Furthermore, our rapid acclimation projections may be unrealistically optimistic because they ignore the potential for transient dynamics involving dispersal limitation (Urban *et al*. 2012) or novel, alternative stable states (Carpenter 2002; Blonder *et al*. 2017) to prevent ecosystem functioning from perfectly tracking a rapidly changing climate.

An improved understanding of ecosystem acclimation rates and processes would be extremely valuable for climate adaptation management. For example, the “Resist-Accept-Direct” (RAD) framework (Lynch *et al*. 2021; Thompson *et al*. 2021; Schuurman *et al*. 2022) provides managers with three management pathways: Resist the trajectory of change, Accept it, or Direct it by actively steering the ecosystem toward a preferred, new configuration. However, managers often lack robust scientific guidance to inform these choices. Projections like ours could help fill this gap. For the California grasslands, both the minimal and rapid acclimation scenarios projected forage increase for most locations, making Accept an easy choice. In every other ecoregion, a Direct strategy to facilitate ecosystem acclimation would help limit production declines, as in the Shortgrass Steppe and Hot Deserts, or even prevent production declines, as in much of the Cold Desert and Northern Mixed Prairie Ecosystems. Of course, forage production is just one ecosystem service, and RAD decisions would consider many services, including biodiversity. In practice, tension may emerge between conserving historical species composition and maintaining ecosystem functioning by promoting compositional change (Fisichelli *et al*. 2016).

Currently, the science is not mature enough to guide climate adaptation management. New research to better understand the rates and impacts of acclimation processes (Fig. 1) is needed to help managers effectively facilitate ecosystem acclimation. Such research would also reduce uncertainty in long-term projections of ecosystem services and advance our understanding of processes linking rapid environmental change, ecosystem structure, and ecosystem function.

## Acknowledgments

This research was supported by the USDA National Institute of Food and Agriculture, Agricultural and Food Research Initiative Competitive Program, Climate and Land Use (grant no. 2018-68002-27923), and by the Utah Agriculture Experiment Station (journal paper #9505). RKS was supported by a Postdoctoral Fellowship (grant no. 2019-67012-29726/project accession no. 1019364) from the USDA National Institute of Food and Agriculture. Any use of trade, firm, or product name is for descriptive purposes only and does not imply endorsement by the U.S. Government. We thank Brady Allred for helping us access the productivity data, and Jonathan Levine, Janneke HilleRisLambers, and Alan Knapp for comments that improved an earlier version of the manuscript.

## Notes

### Competing Interest Statement

The authors have declared no competing interest.

### Summary of Updates

We revised to address reviewer comments on the previous version. We also added a new Figure 1 to explain the concepts.

