## Supporting Information for "The rate of ecosystem acclimation is the dominant uncertainty in long-term projections of an ecosystem service"

**This PDF file includes:**

Supporting Methods
Figures S1 to S9
Tables S1 to S5
Supporting References

### Supporting Methods

#### *Comparison to field data*

The herbaceous production "data" that is our response variable is itself the output of a model. Robinson et al. (2019) derived productivity estimates from remotely sensed data and climate variables, including daily temperature. Our analysis showed high sensitivity of production to growing season mean temperature in many ecoregions. Could that result be an artifact of the way daily temperature is used in Robinson et al.'s (2019) algorithm?

We tested for such an artifact by comparing the temperature sensitivities of production observed in our analysis of the remotely sensed data with temperature sensitivities estimated directly from field data. We found five grassland sites located within our study region for which at least 29 years of aboveground annual production data were available. We focused on grasslands and excluded shrublands to minimize the difference between our remotely sensed data on herbaceous production and the total aboveground production data collected in the field. The five sites included the San Joaquin Experimental Range in the California Annual grasslands ecoregion, the Central Plains Experimental Range in the Shortgrass Steppe ecoregion, and three grassland sites at the Jornada Experimental Range, in the Hot Deserts summer precipitation ecoregion.

For each of these sites, we calculated the relative (proportional) sensitivity of production to precipitation three different ways: 1) We regressed the field production data on annual growing season precipitation and temperature from PRISM (https://prism.oregonstate.edu/) for the field site location. We defined growing seasons for each ecoregion consistent with our analysis of the remotely sensed data. We then calculated the relative change in predicted production between a hypothetical year with mean precipitation and temperature, and a hypothetical year with mean precipitation but temperature 1°C above the mean. 2) We regressed the remotely sensed data (for the pixel containing the field location) on annual growing season precipitation and temperature data from PRISM, rather than on annual weather data from the Livneh et al. (2015) data set we used in our primary analysis. We then calculated temperature sensitivity exactly as we did for the field data set. 3) We calculated temperature sensitivity for the field site location based on our regional-scale analysis of the remotely-sensed production data and the Livneh et al. (2015) historical weather data.

The results show no evidence of a clear bias towards higher temperature sensitivities when remotely-sensed production is the response variable (Table S5). The results did not change when we included an interaction between precipitation and temperature in our local-scale analyses.

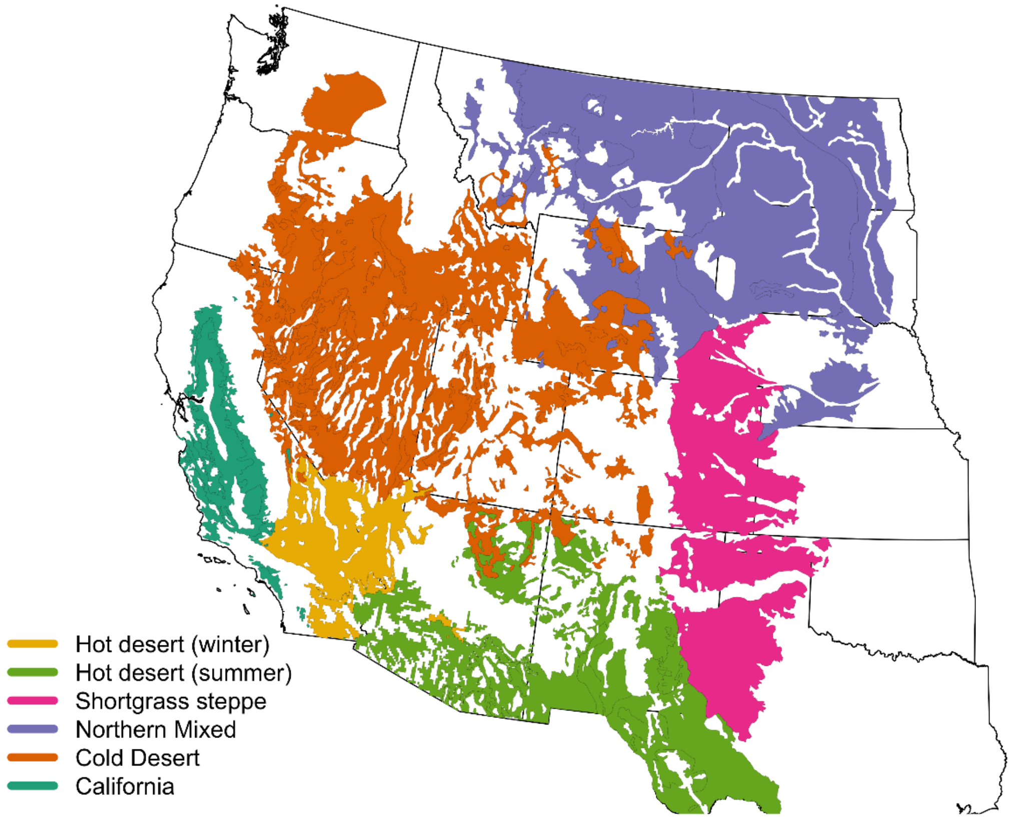


**Fig. S1.** Map of the six rangeland ecoregions we studied. The ecoregions were initially based on a map of potential natural vegetation (Kuchler 1964).

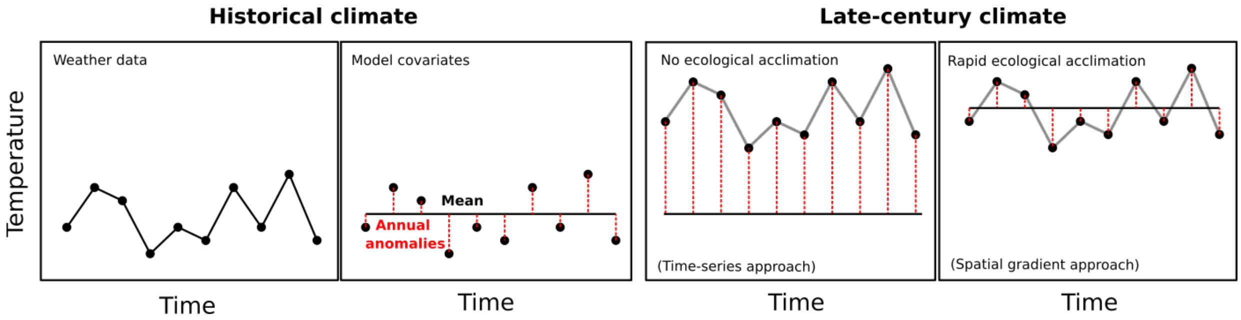


**Fig. S2.** The difference between the time-series and spatial gradient approaches. The first panel on the left shows a hypothetical, historical time-series of a weather variable, temperature in this case, for one location. The second panel shows how we convert this time-series into two model covariates, a long-term mean, the values of which vary only in space, from one location to another, and annual anomalies, which vary only through time at each location. To project future forage production, we use future weather data simulated by a general circulation model (GCM). The grey lines in the two panels on the right show a future temperature time-series for our hypothetical location. We can decompose the future temperature series into spatial and temporal components in two different ways. First, we can calculate the annual anomalies based on the historic mean (third panel). This approach results in no change in the mean covariates but large changes in the annual anomalies. We call this the time-series approach, and it assumes minimal ecological acclimation. Second, we can calculate the annual anomalies based on the future means (fourth panel). This approach results in a large change in the mean but very little change in the annual anomalies. We call this the spatial gradient approach, and it assumes very rapid ecological acclimation.

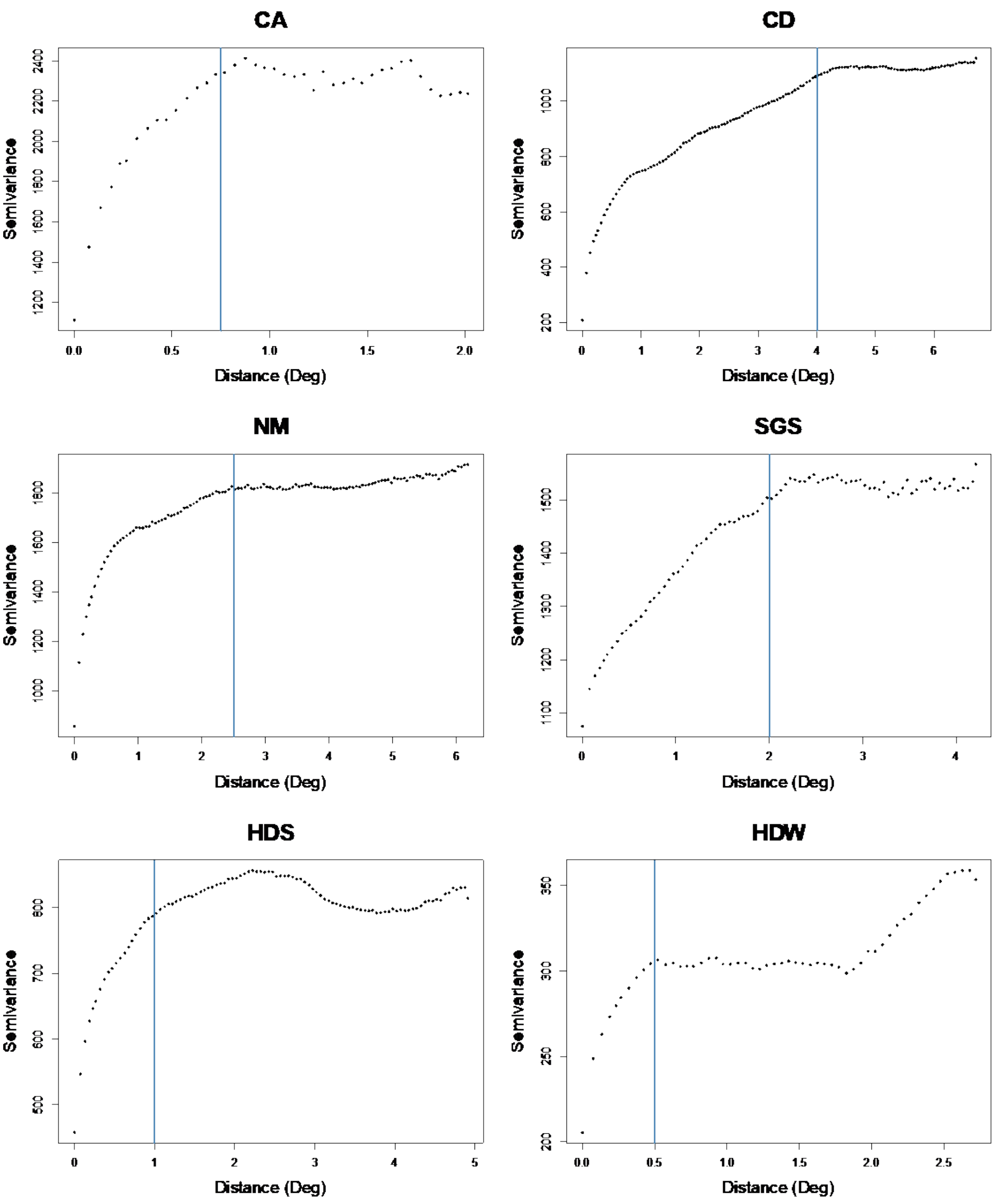


**Fig. S3.** Semivariograms summarizing residual spatial autocorrelation in models after accounting for fixed effects of average climate conditions and annual deviations in each ecoregion. Points indicate semivariance estimates and the vertical blue bar is the autocorrelation range used in fitting the full model.

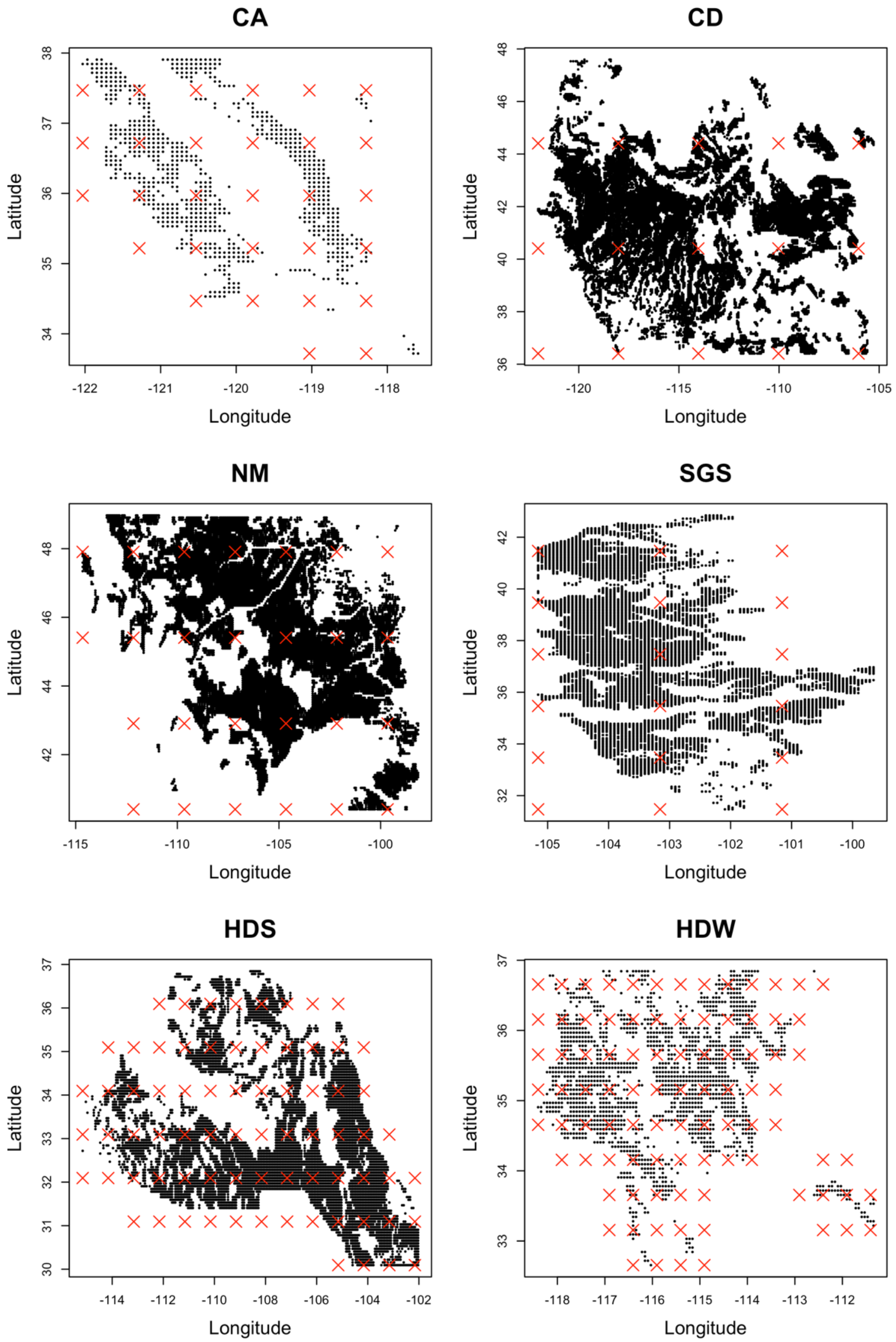


**Fig. S4.** Locations of individual pixels (black points) and knots (red X's) used to model spatial autocorrelation in each ecoregion.

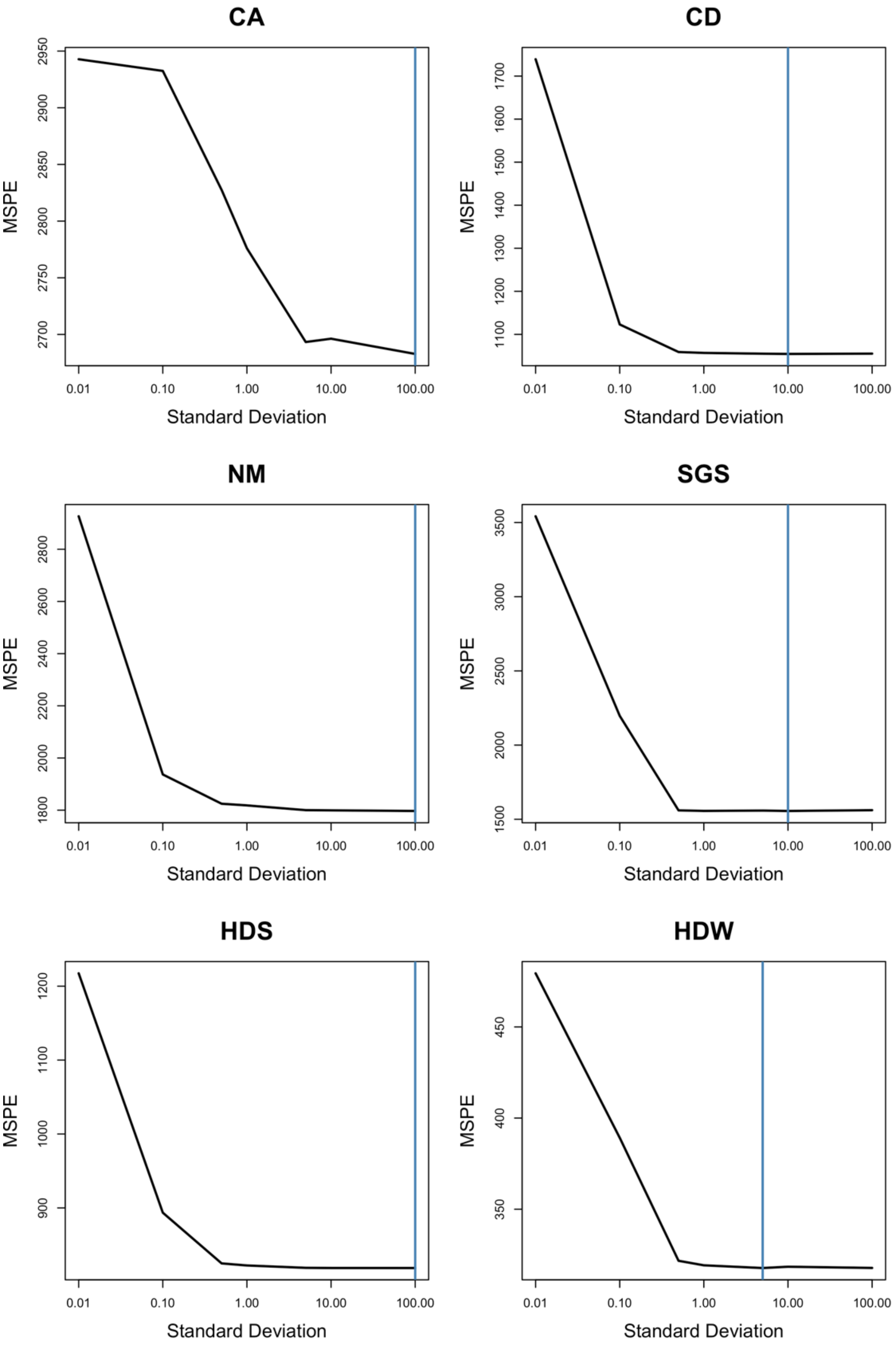


**Fig. S5.** Results of ridge regression analysis to minimize covariate overfitting. The standard deviation of the covariate priors was sequentially reduced to identify the priors that lead to the lowest predictive error (MSPE) with withheld data. The vertical blue bar indicates the prior standard deviation with the smallest predictive error.

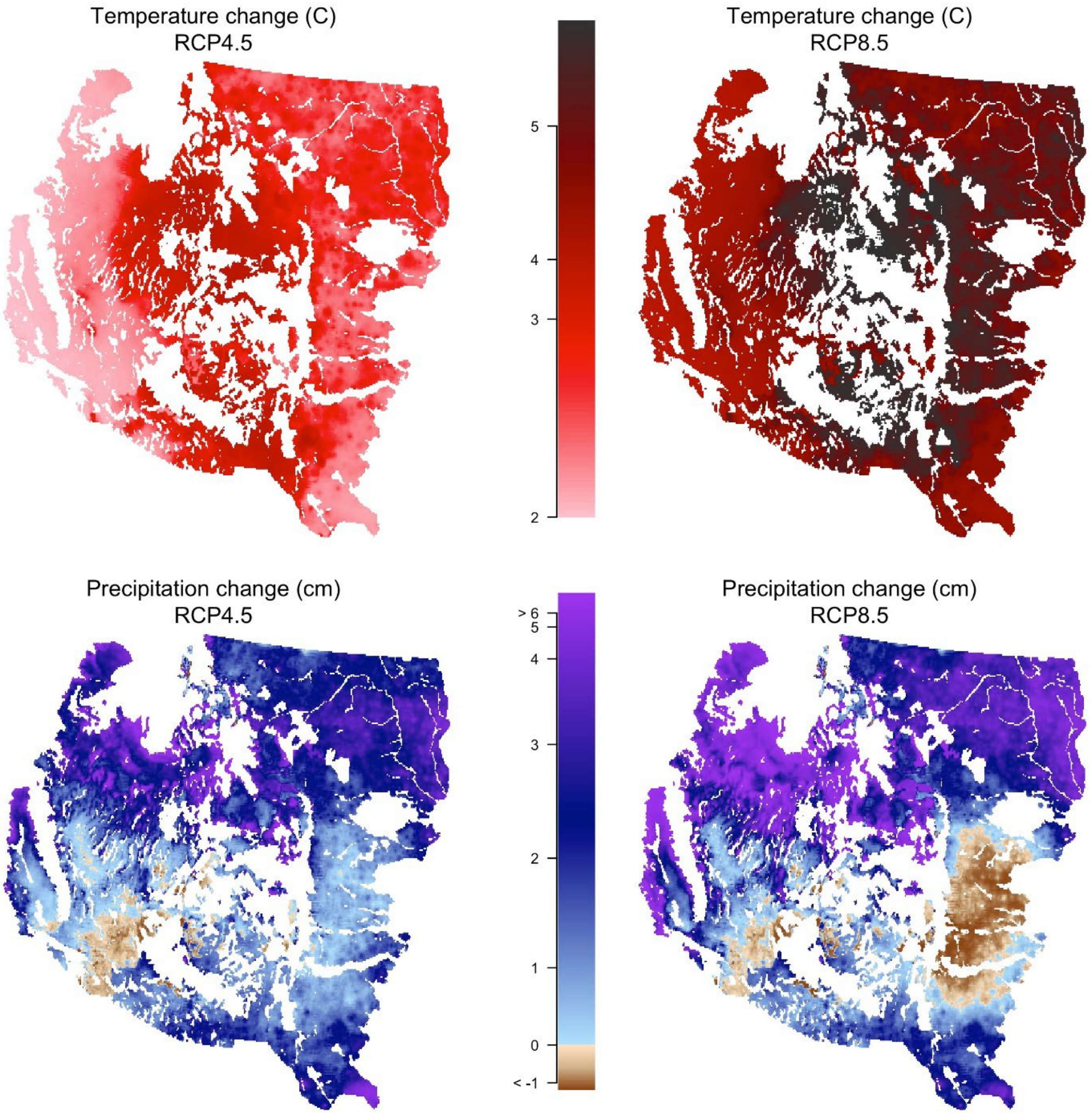


**Fig. S6**. Projected changes in growing season temperature (top) and precipitation (bottom) for RCP 4.5 (left) and 8.5 (right). Values are differences between the late century mean and the historical mean, averaged across the 11 GCMs in our ensemble.

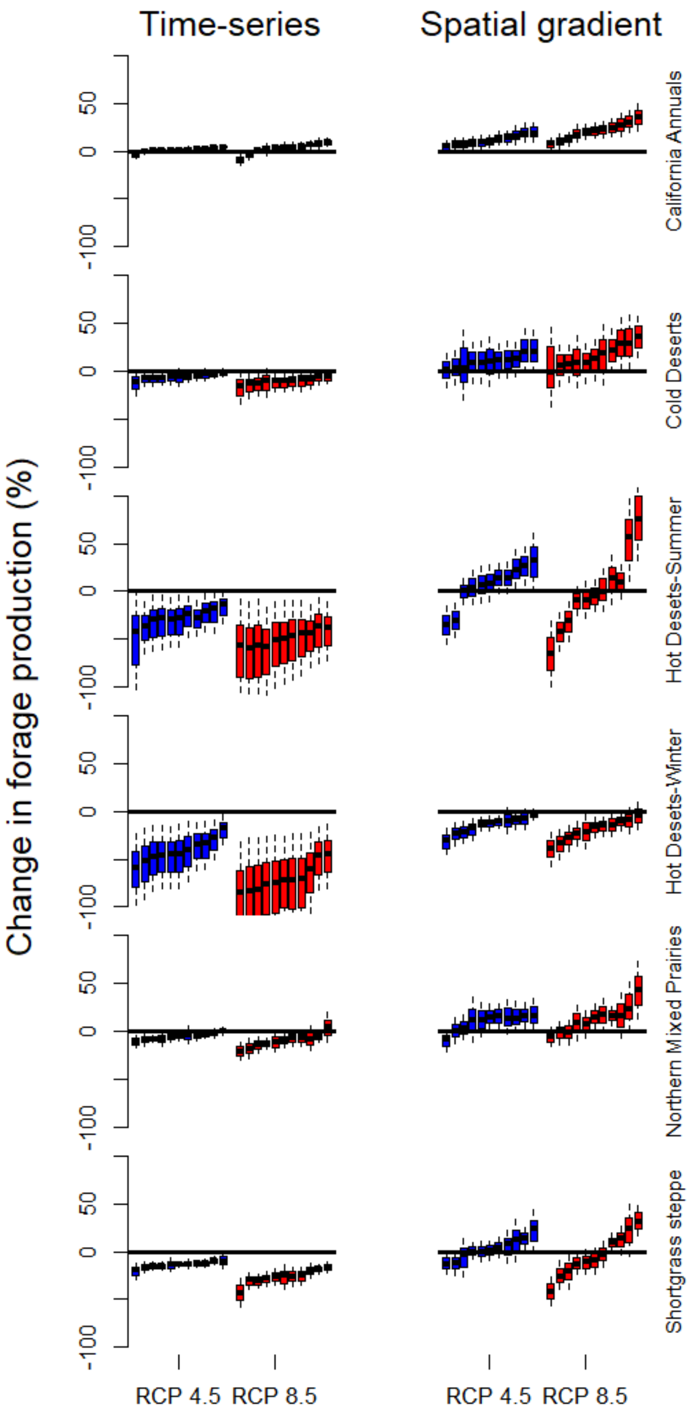


**Fig. S7.** Projected changes in forage production for each combination of the 11 GCMs, two RCPs and two projection methods for each ecoregion (rows). Each box shows one GCM and RCP combination for each ecoregion and projection method. Boxes depict the interquartile range of the data (solid line inside is the median) and whiskers are equal to 0.5 times the interquartile range.

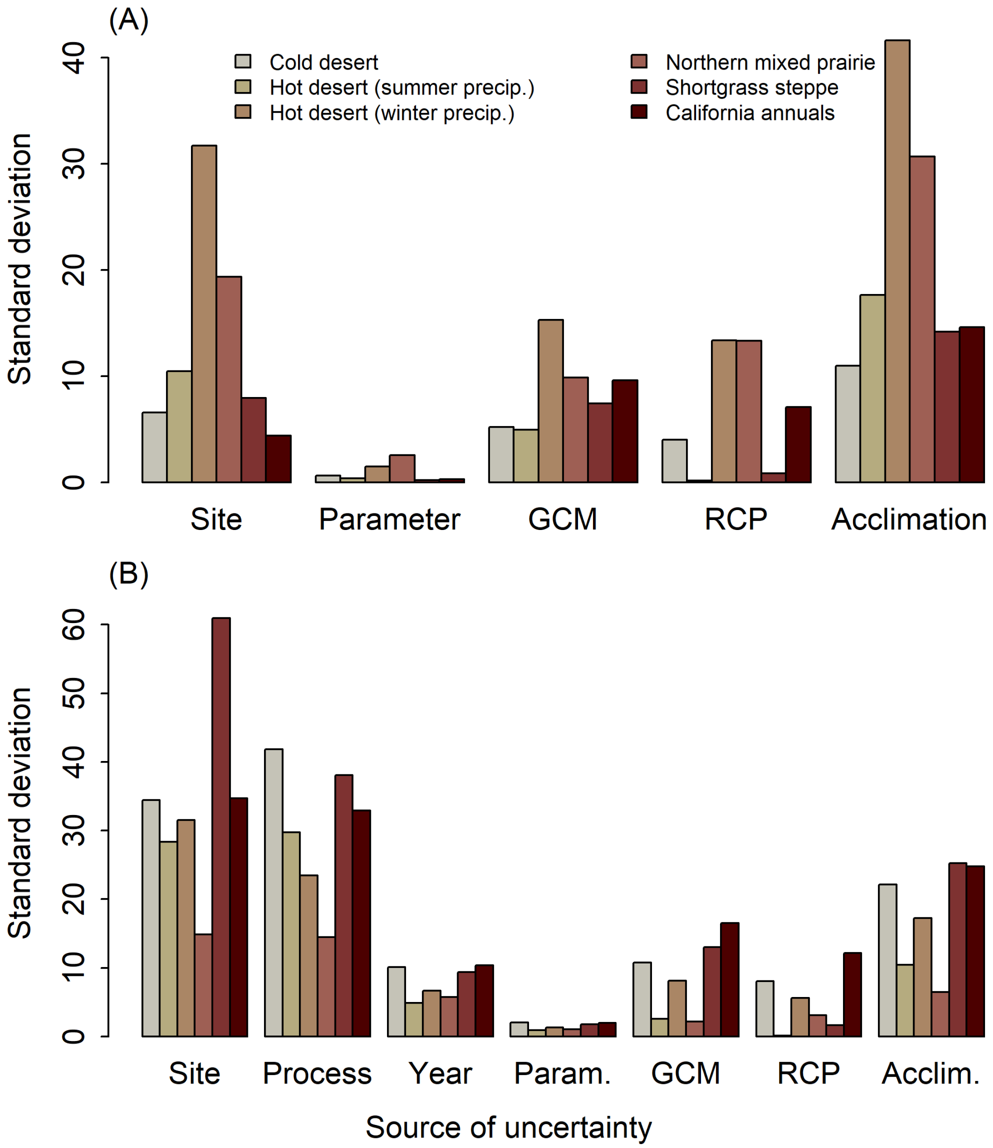


**Fig. S8.** Sources of variation in future forage production, quantified as the standard deviation in the marginal means of the sources of uncertainty. (A) Variation in projected changes in forage production. (B) Variation in absolute forage production projected for the late 21$^{st}$ century. The greater variation among sites, relative to panel A, reflects strong spatial gradients in precipitation and temperature. This analysis also includes uncertainty from process error and from random year effects ("Year"), which do not appear in the analysis of productivity changes because we assume they are equivalent in the historic and future periods. GCM: global circulation model. RCP: relative concentration pathway (greenhouse gas emission scenario). Acclimation: differences in the rate of ecological acclimation captured by the contrast between the time-series and spatial gradient approaches.

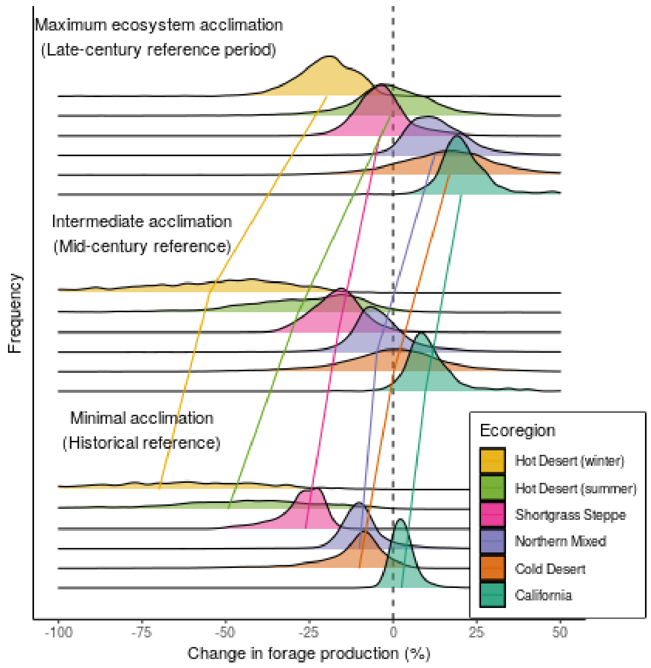


**Fig. S9.** Projected changes in forage production as a function of the rate of ecological acclimation. For each ecoregion (colors), we show the distribution of projected changes in forage production among locations for three scenarios: (top) rapid acclimation, (middle) an intermediate rate of acclimation, and (bottom) minimal acclimation. Lines indicate the median rate of change for each ecoregion and acclimation scenario. The change for each location is an average of projections using 11 GCMs for the RCP 8.5 emission pathway. Locations where decreases in forage production exceeded -100% are not shown, but were included in calculations of medians. "Reference" refers to the time period used to calculate climate means for each location, which were then used to calculate annual anomalies.

**Table S1.** Derived covariates and interactions.

| Covariate | Interpretation |
|---|---|
| $\bar{P}$ | Mean total precipitation, captures spatial variation in productivity explained by spatial variation in precipitation. |
| $\delta\mathrm{P}$ | Annual precipitation anomaly, captures temporal variation in productivity explained by temporal variation in precipitation within a location. |
| $\bar{\mathrm{T}}$ | Mean temperature, captures spatial variation in productivity explained by spatial variation in temperature. |
| $\delta\mathrm{T}$ | Annual temperature anomaly, captures temporal variation in productivity explained by temporal variation in temperature within a location. |
| $\bar{P} \times \delta P$ | This interaction term allows the effect of precipitation anomalies on productivity to vary in space. For example, a wet year might have a weaker effect on productivity at a wet site than at a dry site. |
| $\delta\mathrm{P}^2$ | This interaction allows for a saturating (nonlinear) relationship between productivity and precipitation anomalies. |
| $\bar{\mathrm{P}} \times \delta\mathrm{P}^2$ | This interaction allows the saturating effect of precipitation anomalies to vary across space with mean precipitation. |
| $\bar{T} \times \delta T$ | This interaction term allows the effect of temperature anomalies on productivity to vary in space. For example, a hot year might have a positive effect on productivity at a cold site but a negative effect at a hot site. |
| $\delta\mathrm{P} \times \delta\mathrm{T}$ | This interaction allows the effect of wet year on productivity to depend on the temperature in that year. |

**Table S2.** Parameters estimates for each of the six ecoregion-specific models.

| Parameter | Mean | 2.5% CI | 97.5% CI |
|---|---|---|---|
| **CA** | | | |
| $Intercept\ (b_1)$ | 206.85 | 203.07 | 210.48 |
| $\bar{P}\ (b_2)$ | 26.55 | 25.81 | 27.27 |
| $\delta\text{P}\ (b_3)$ | 22.97 | 19.45 | 26.5 |
| $\bar{\text{T}}\ (b_4)$ | 15.48 | 14.86 | 16.1 |
| $\delta\text{T}\ (b_5)$ | 3.22 | -0.7 | 7.13 |
| $\bar{P} \times \delta P\ (b_6)$ | -15.83 | -18.53 | -13.23 |
| $\delta\text{P}^2\ (b_7)$ | -19.31 | -22.15 | -16.37 |
| $\bar{P} \times \delta\text{P}^2\ (b_8)$ | 13.16 | 10.55 | 15.74 |
| $\bar{T} \times \delta T\ (b_9)$ | -2.73 | -6.49 | 1.08 |
| $\delta\text{P} \times \delta T\ (b_{10})$ | 3.73 | 2.86 | 4.62 |
| $\sigma_{(s)}$ | 36.09 | 34.41 | 37.78 |
| $\sigma_{(p)}$ | 41.81 | 41.43 | 42.2 |
| $\sigma_{(t)}$ | 10.12 | 9.21 | 11.11 |
| **CD** | | | |
| $Intercept\ (b_1)$ | 62.17 | 60.21 | 63.85 |
| $\bar{P}\ (b_2)$ | 26.44 | 26.32 | 26.56 |
| $\delta\text{P}\ (b_3)$ | 7.25 | 6.94 | 7.54 |
| $\bar{\text{T}}\ (b_4)$ | -0.02 | -0.16 | 0.12 |
| $\delta\text{T}\ (b_5)$ | -0.39 | -0.6 | -0.19 |
| $\bar{P} \times \delta P\ (b_6)$ | -2.08 | -2.34 | -1.82 |
| $\delta\text{P}^2\ (b_7)$ | 2.99 | 2.67 | 3.3 |
| $\bar{P} \times \delta\text{P}^2\ (b_8)$ | -7.92 | -8.22 | -7.62 |
| $\bar{T} \times \delta T\ (b_9)$ | -1.05 | -1.22 | -0.89 |
| $\delta\text{P} \times \delta T\ (b_{10})$ | -0.35 | -0.46 | -0.24 |
| $\sigma_{(s)}$ | 163.22 | 161.11 | 165.34 |
| $\sigma_{(p)}$ | 29.76 | 29.68 | 29.83 |
| $\sigma_{(t)}$ | 4.89 | 4.19 | 5.73 |
| **HDS** | | | |
| $Intercept\ (b_1)$ | 55.5 | 53.11 | 58.17 |
| $\bar{P}\ (b_2)$ | 25.38 | 25.15 | 25.61 |
| $\delta\text{P}\ (b_3)$ | 1.49 | 0.98 | 2.01 |
| $\bar{\text{T}}\ (b_4)$ | -3.71 | -3.97 | -3.46 |
| $\delta\text{T}\ (b_5)$ | 8.04 | 7.24 | 8.81 |
| $\bar{P} \times \delta P\ (b_6)$ | 7.97 | 7.49 | 8.44 |
| $\delta\text{P}^2\ (b_7)$ | -3.11 | -3.67 | -2.54 |
| $\bar{P} \times \delta\text{P}^2\ (b_8)$ | 0.73 | 0.15 | 1.3 |
| $\bar{T} \times \delta T\ (b_9)$ | -12.31 | -13.06 | -11.53 |
| $\delta\text{P} \times \delta T\ (b_{10})$ | -1.33 | -1.49 | -1.17 |
| $\sigma_{(s)}$ | 104.49 | 103.29 | 105.68 |
| $\sigma_{(p)}$ | 23.45 | 23.38 | 23.53 |
| $\sigma_{(t)}$ | 6.66 | 5.85 | 7.56 |

| | | | |
|---|---|---|---|
| **HDW** | | | |
| $Intercept\ (b_1)$ | 24.14 | 22.12 | 26.12 |
| $\bar{P}\ (b_2)$ | 3.84 | 3.54 | 4.15 |
| $\delta P\ (b_3)$ | 7.19 | 6.37 | 8.04 |
| $\bar{T}\ (b_4)$ | -1.9 | -2.23 | -1.59 |
| $\delta T\ (b_5)$ | 2.46 | 1.62 | 3.3 |
| $\bar{P} \times \delta P\ (b_6)$ | 0.94 | 0.3 | 1.59 |
| $\delta P^2\ (b_7)$ | 1.92 | 1.21 | 2.62 |
| $\bar{P} \times \delta P^2\ (b_8)$ | -4.53 | -5.13 | -3.9 |
| $\bar{T} \times \delta T\ (b_9)$ | -5.55 | -6.36 | -4.72 |
| $\delta P \times \delta T\ (b_{10})$ | -1.23 | -1.46 | -1 |
| $\sigma_{(s)}$ | 43.82 | 42.67 | 44.97 |
| $\sigma_{(p)}$ | 14.46 | 14.36 | 14.55 |
| $\sigma_{(t)}$ | 5.73 | 4.96 | 6.58 |
| **NM** | | | |
| $Intercept\ (b_1)$ | 197.01 | 193.67 | 200.36 |
| $\bar{P}\ (b_2)$ | 64.17 | 63.99 | 64.34 |
| $\delta P\ (b_3)$ | 16.39 | 15.74 | 17.04 |
| $\bar{T}\ (b_4)$ | 1.05 | 0.91 | 1.2 |
| $\delta T\ (b_5)$ | 15.91 | 14.66 | 17.19 |
| $\bar{P} \times \delta P\ (b_6)$ | 0.07 | -0.54 | 0.68 |
| $\delta P^2\ (b_7)$ | 5.95 | 5.36 | 6.54 |
| $\bar{P} \times \delta P^2\ (b_8)$ | -7.51 | -8.1 | -6.91 |
| $\bar{T} \times \delta T\ (b_9)$ | -21.54 | -22.77 | -20.33 |
| $\delta P \times \delta T\ (b_{10})$ | 3.88 | 3.68 | 4.07 |
| $\sigma_{(s)}$ | 114.76 | 112.8 | 116.7 |
| $\sigma_{(p)}$ | 38.05 | 37.96 | 38.14 |
| $\sigma_{(t)}$ | 9.37 | 8.5 | 10.32 |
| **SGS** | | | |
| $Intercept\ (b_1)$ | 174.62 | 171.16 | 178.34 |
| $\bar{P}\ (b_2)$ | 36.38 | 36.19 | 36.57 |
| $\delta P\ (b_3)$ | 27.04 | 25.71 | 28.34 |
| $\bar{T}\ (b_4)$ | -1.26 | -1.43 | -1.08 |
| $\delta T\ (b_5)$ | -0.13 | -1.53 | 1.27 |
| $\bar{P} \times \delta P\ (b_6)$ | -10.87 | -12.14 | -9.6 |
| $\delta P^2\ (b_7)$ | 2.73 | 1.42 | 4 |
| $\bar{P} \times \delta P^2\ (b_8)$ | -5.3 | -6.56 | -4.04 |
| $\bar{T} \times \delta T\ (b_9)$ | -7.25 | -8.65 | -5.88 |
| $\delta P \times \delta T\ (b_{10})$ | 1.06 | 0.76 | 1.37 |
| $\sigma_{(s)}$ | 1.22 | 0.05 | 3.39 |
| $\sigma_{(p)}$ | 32.93 | 32.81 | 33.05 |
| $\sigma_{(t)}$ | 10.34 | 9.48 | 11.3 |

**Table S3.** Goodness-of-fit measures for the six ecoregion-specific statistical model. Values are squared correlation coefficients for all observations and predictions ("All"), the observed and predicted means (averaged across years) for each location ("Spatial"), and the observed and predicted annual forage anomalies ("Temporal"). Predictions were calculated using fixed effects only.

| Ecoregion | All | Spatial | Temporal |
|---|---|---|---|
| CA | 0.24 | 0.38 | 0.37 |
| CD | 0.39 | 0.44 | 0.45 |
| HDS | 0.5 | 0.64 | 0.52 |
| HDW | 0.32 | 0.18 | 0.28 |
| NM | 0.71 | 0.8 | 0.74 |
| SGS | 0.54 | 0.73 | 0.61 |

**Table S4.** Comparison of sensitivity of production to temperature based on field-based and remotely-sensed estimates of production. Sensitivity is expressed as the proportional change in production caused by a 1°C increase in temperature. Methods are described in Supporting Methods.

| **Site** | **Ecoregion** | **Field-based** | **Remotely-sensed, site-specific model** | **Remotely-sensed, regional model** |
|---|---|---|---|---|
| Jornada Experimental Range, BASN site | HDS | -0.18 | -0.25 | -0.10 |
| Jornada Experimental Range ,IBPE site | HDS | -0.05 | -0.21 | -0.10 |
| Jornada Experimental Range, SUMM site | HDS | -0.03 | -0.10 | -0.09 |
| Central Plains Experimental Range | SGS | -0.16 | -0.09 | -0.06 |
| San Joaquin Experimental Range | CA | -0.05 | -0.01 | 0.00 |

**Table S5.** Sum-of-squares components for each of the six ecoregions. Values are fractions of the total sum-of-squares.

| Source | CA | CD | HDS | HDW | NM | SGS |
|---|---|---|---|---|---|---|
| Site | 0.246 | 0.21 | 0.267 | 0.273 | 0.18 | 0.052 |
| Parameter error | 0.002 | <0.0005 | 0.001 | 0.005 | <0.0005 | <0.0005 |
| GCM | 0.141 | 0.043 | 0.057 | 0.065 | 0.143 | 0.226 |
| RCP | 0.046 | <0.0005 | 0.024 | 0.064 | 0.001 | 0.068 |
| Acclimation rate | 0.342 | 0.299 | 0.23 | 0.342 | 0.286 | 0.287 |
| Acclim. rate × Site | 0.085 | 0.193 | 0.228 | 0.121 | 0.109 | 0.058 |
| Acclim. rate × Param. error | 0.002 | <0.0005 | 0.001 | 0.002 | <0.0005 | <0.0005 |
| Acclim. rate × GCM | 0.023 | 0.027 | 0.072 | 0.018 | 0.036 | 0.094 |
| Acclim. rate × RCP | 0.023 | 0.017 | 0.01 | 0.033 | 0.009 | 0.008 |
| Sum of all Acclim. ×'s | 0.133 | 0.238 | 0.311 | 0.175 | 0.154 | 0.16 |
| Sum of all Acclim. terms | 0.475 | 0.537 | 0.541 | 0.517 | 0.441 | 0.446 |
| Sum of all computed components | 0.909 | 0.791 | 0.889 | 0.924 | 0.765 | 0.792 |
